# Facilitative priority effects drive parasite assembly under coinfection

**DOI:** 10.1101/2020.03.30.015495

**Authors:** Fletcher W. Halliday, Rachel M. Penczykowski, Benoit Barrès, Jenalle L. Eck, Elina Numminen, Anna-Liisa Laine

## Abstract

Host individuals are often coinfected with diverse parasite assemblages, resulting in complex interactions among parasites within hosts. Within hosts, priority effects occur when the infection sequence alters the outcome of interactions among parasites. Yet, the role of host immunity in this process remains poorly understood. We hypothesized that the host response to first infection could generate priority effects among parasites, altering the assembly of later arriving strains during epidemics. We tested this by infecting sentinel host genotypes of *Plantago lanceolata* with strains of the fungal parasite, *Podosphaera plantaginis*, and measuring susceptibility to subsequent infection during experimental and natural epidemics. In these experiments, prior infection by one strain often increased susceptibility to other strains, and these facilitative priority effects altered the structure of parasite assemblages, but this effect depended on host genotype, host population, and parasite genotype. Thus, host genotype, spatial structure, and priority effects among strains all independently altered parasite assembly. Then, using a fine-scale survey and sampling of infections on wild hosts in several populations, we identified a signal of facilitative priority effects, which altered parasite assembly during natural epidemics. Together, these results provide evidence that within host priority effects by early arriving strains can drive parasite assembly, with implications for how strain diversity is spatially and temporally distributed during epidemics.

## Introduction

The diversity of parasites – organisms that live in and on hosts, potentially causing disease – may rival the diversity of all other organisms on earth (Dobson *et al.* 2008). In light of this diversity, it is not surprising that host individuals are often infected with diverse parasite assemblages, composed of multiple parasite species or multiple genetic variants (‘strains’) of the same species (Mideo 2009; Greischar *et al.* 2020). Within hosts, interactions among coinfecting parasite strains can influence the dynamics of drug resistance (Wale *et al.* 2017), evolution of virulence (Bhattacharya *et al.* 2019), and the magnitude of parasite epidemics (Susi *et al.* 2015), with implications for host health (Read & Taylor 2001). Thus, understanding how parasite strains interact in shared host individuals may be important for predicting the spread of infectious diseases and ameliorating their impact on host populations. Yet, measuring how interactions among parasites influence natural epidemics is notoriously difficult, as this requires manipulating focal mechanisms of interactions and documenting the structure of parasite assemblages as epidemics unfold (Mideo 2009; Hawley & Altizer 2011; Hoverman *et al.* 2013; Zhan & McDonald 2013; Hellard *et al.* 2015; Tollenaere *et al.* 2015; Budischak *et al.* 2018). Using a parasitic fungus that infects a wild host plant, this study experimentally tests whether parasite interactions that are mediated by the host response to initial infection alter the structure of parasite assemblages within hosts under field conditions, and then leverages the results of these experiments to explain how parasite strains assemble during a natural epidemic.

Multiple parasites that encounter the same host individual can interact during simultaneous infections, known as coinfections (Griffiths *et al.* 2014; Tollenaere *et al.* 2015; Ezenwa 2016). One potential mechanism of interaction among coinfecting parasites occurs when host immune responses to one parasite alter host susceptibility to secondary infections of another parasite (Lello *et al.* 2004; Mideo 2009; Chung *et al.* 2012; Halliday *et al.* 2018). This mechanism can result in either antagonism or facilitation among coinfecting parasites, and ultimately can alter parasite epidemics (Eswarappa *et al.* 2012; Tollenaere *et al.* 2015; Zélé *et al.* 2018). The immune response to initial infection can suppress coinfection when infection by one parasite activates immune signaling pathways that induce resistance to subsequent infections, in a process known by a variety of terms including immune priming, cross protection, induced resistance, or cross-immunity (Jenner 1923; Fulton 1986; Van Loon 1997; Conrath *et al.* 2006; Pieterse *et al.* 2014). Alternatively, an early arriving parasite can facilitate coinfection by inactivating immune signaling pathways that protect hosts from multiple parasites (Spoel *et al.* 2007; Kliebenstein & Rowe 2008). These effects can be temporary and spatially restricted within hosts (Koornneef *et al.* 2008), or systemic and persistent long after initial infection (Pieterse *et al.* 2014). Both mechanisms of immune-mediated interactions among parasites have been reported in plant and animal hosts (Glazebrook 2005; Ezenwa *et al.* 2010; Pieterse *et al.* 2014). These effects, which have been predominantly tested in laboratory environments (but see Halliday *et al.* 2018), indicate that the sequence and timing of infections may influence the structure of parasite assemblages.

The field of community ecology provides a framework for understanding how the sequence of infection on host individuals might alter parasite assemblages as epidemics unfold (Hoverman *et al.* 2013; Vannette & Fukami 2014; Fukami 2015; Johnson *et al.* 2015; Halliday *et al.* 2017; Clay *et al.* 2019; Karvonen *et al.* 2019). Specifically, interactions among parasites that are contingent on the sequence of past events can be a consequence of priority effects within hosts. Within hosts, priority effects occur when the per-capita strength of antagonism or facilitation among parasites is altered by their sequence of arrival (Hoverman *et al.* 2013; Mordecai *et al.* 2016). Priority effects, in turn can drive community assembly, thereby altering the structure of parasite communities during natural epidemics (Halliday *et al.* 2017; Clay *et al.* 2020). Priority effects are expected to occur most commonly when species exhibit high niche overlap and when early arriving species have large impacts on the availability of that niche (Vannette & Fukami 2014). A host comprises the entire niche available to parasites during infection (Kuris *et al.* 1980; Rynkiewicz *et al.* 2015), and thus coinfecting parasites often exhibit high niche overlap (Sousa 1992; Graham 2008; Seabloom *et al.* 2015), particularly when parasite assemblages are comprised of coinfecting strains of the same parasite species (e.g., Wale *et al.* 2017). Although priority effects have been predominantly used to describe community assembly in multi-species parasite assemblages (reviewed in Clay *et al.* 2019), these same principles may apply to parasite assemblages comprised of multiple strains (Greischar *et al.* 2020). By activating immune responses that alter host susceptibility, early arriving strains can therefore determine the availability of the shared host niche (Cobey & Lipsitch 2013); thus, the immune response to initial infection may drive priority effects among parasite strains within hosts, thereby altering the structure of parasite assemblages within hosts.

The degree to which the sequence and timing of infection influences parasite assemblages might depend on the history of interactions between host and parasite populations (Tollenaere *et al.* 2015). This history of interactions between host and parasite populations, which is typically measured through local adaptation assays (Greischar & Koskella 2007; Hoeksema & Forde 2008), is commonly reflected by differences in the susceptibility of certain host genotypes to certain parasite genotypes (Burdon & Laine 2019). Interactions among sequentially arriving parasites could also depend on host or parasite genotypes if a given host genotype is more or less sensitive to infection by the first or second arriving parasite genotype (Lambrechts *et al.* 2006), or if the response triggered by the first arriving parasite is genotype specific (Ferro *et al.* 2019; Westman *et al.* 2019). Thus, whether or not within-host priority effects alter parasite epidemics might depend on complex interactions among host and parasite genotypes. Consequently, it is essential to incorporate genotypic variation into studies of sequential infection among parasite strains.

The host response to infection may alter parasite interactions and epidemics (Lello *et al.* 2004; Graham 2008; Tollenaere *et al.* 2015), but we lack studies that experimentally manipulate prior parasite exposure and measure the consequences for parasites outside of the lab (Hellard *et al.* 2015; Pedersen & Fenton 2015; Budischak *et al.* 2018), hampering our understanding of the general processes through which within-host parasite interactions alter parasite assemblages in nature. This study addresses this research gap experimentally by first infecting host plants with parasitic fungi, physically restricting those parasites from interacting directly within hosts, and then testing whether the host response to initial infection alters the structure of parasite assemblages. We then leverage the experimental results to explain how parasite strains assemble within hosts during a natural epidemic. We find that parasites exhibit facilitative priority effects driven by the host response to initial infection, and that these facilitative priority effects can alter the structure of parasite assemblages during a natural epidemic. These results indicate that the sequence of infection can determine the probability of coinfection, altering the trajectory of parasite assembly, and leading to pronounced differences in the structure of parasite assemblages among hosts.

## Results & Discussion

In order to examine the role of priority effects among parasites that are mediated by the host (i.e., plant) in response to prior infection and the influence of priority effects on the structure of parasite assemblages, we carried out two experiments, referred to as the “manipulated epidemic experiment” and the “natural epidemic experiment”, and a fine-scale survey and sampling of infections in the wild, referred to as the “wild host survey”, using the focal host *Plantago lanceolata*, and the obligate parasite *Podosphaera plantaginis* (Fig 1).

**Figure 1.**
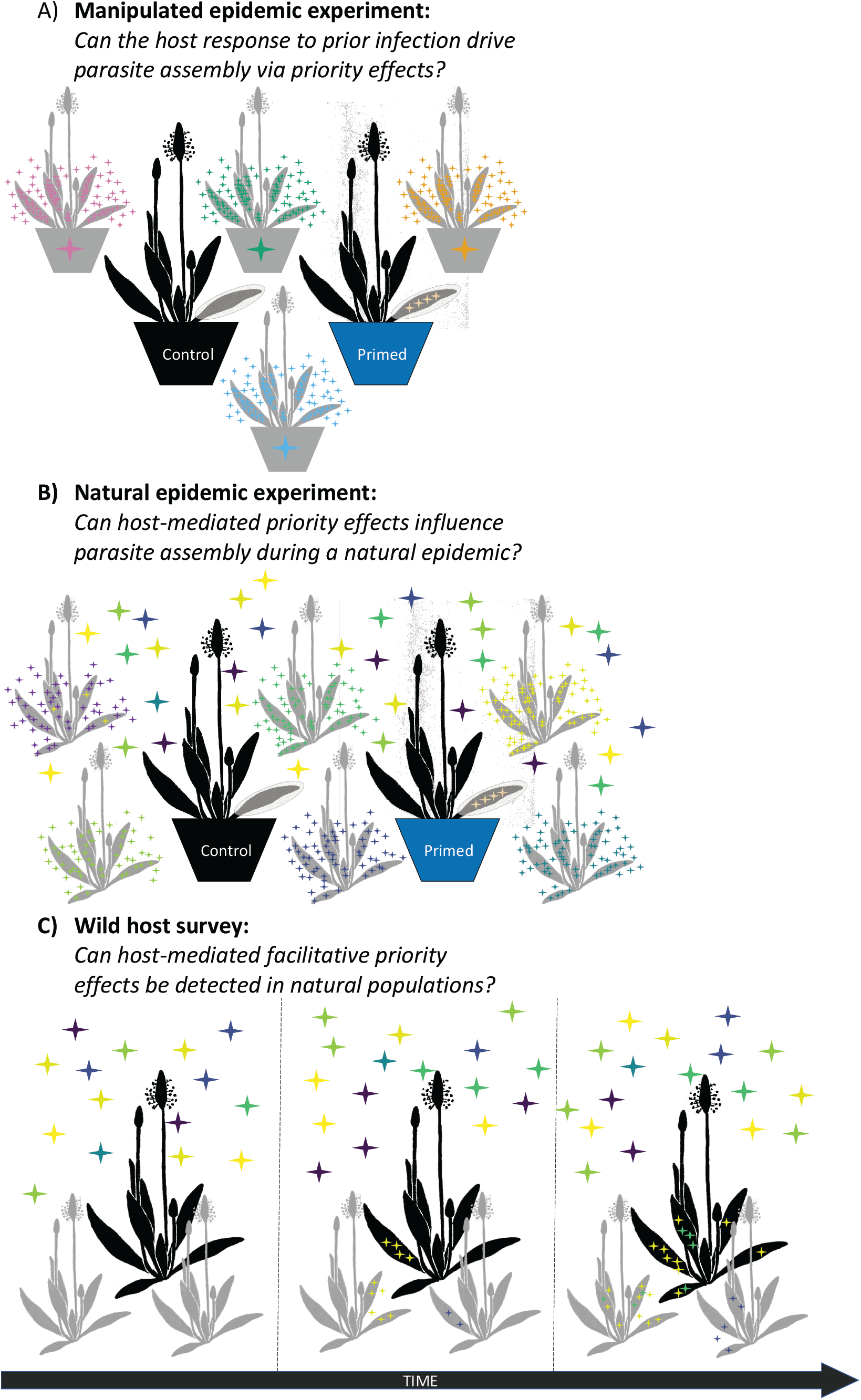
Illustration of key differences and similarities among the three field studies presented in this manuscript. Pot color represents different priming treatments (black = control; blue = primed). Star color represents different parasite genotypes (i.e., strains). A) To test whether host (i.e., plant) responses to prior infection can drive parasite assembly via priority effects, focal potted hosts (black) were either inoculated or mock-inoculated with one of four priming strains. Then hosts were exposed to all four priming strains by placing heavily infected potted hosts (grey) adjacent to the focal hosts under field conditions. This experiment included four different host genotypes not depicted in this figure. B) To test whether host-mediated priority effects among parasites can influence parasite assembly during a natural epidemic, focal potted hosts (black) were either inoculated or mock inoculated with a priming strain associated with a given host population and then embedded in a wild host population (grey) during a natural epidemic. This experiment included four different host genotypes, two infection timing treatments, and three host populations that are not depicted in this figure. C) To test whether a signal of host-mediated facilitative priority effects among parasites could be detected in natural populations, focal wild hosts (black) occurring in wild host populations (grey) were repeatedly surveyed over time. This experiment included 13 host populations that are not depicted in this figure.

### Can host responses to prior infection drive parasite assembly via priority effects?

We carried out the manipulated epidemic experiment in a common garden at the Lammi Biological Station to test whether parasite strains exhibit priority effects that are mediated by the host response to initial infection (Fig. 1). Both the host and parasite species naturally occur in this location. In the manipulated epidemic experiment, four host genotypes were either inoculated or mock-inoculated with one of four parasite strains, which were sealed inside mesh pollination bags to prevent direct strain interactions, and then exposed to all four priming strains for four days. To be consistent with previously published literature (Laine 2011; Conrath *et al.* 2015; Douma *et al.* 2017; Mauch-Mani *et al.* 2017), we refer to the experimental treatment as the “priming treatment” and the experimentally inoculated strains as “priming strains” (Fig. 1).

One challenge of predicting how within-host interactions will alter infection outcomes during epidemics is the difficulty of isolating host-mediated interactions from other interactions among parasites, such as resource or interference competition (Mideo 2009; Budischak *et al.* 2015, 2018). We overcame this limitation experimentally by leveraging the modular growth form of plant hosts. Specifically, for foliar parasites in plant hosts, resource and interference competition are expected to be strongest within individual host leaves (Tollenaere *et al.* 2015; Borer *et al.* 2016; Halliday *et al.* 2017). Because powdery mildews only feed within individual host leaves (Bushnell 2002) and the priming strain was restricted from spreading beyond the inoculated host leaf onto the rest of the host plant, any response to experimental inoculation can be interpreted as an effect that is mediated by the host response to initial infection. Thus, the inoculation treatment was intended to test whether initial infection by one parasite could “prime” the host to respond differently upon subsequent exposure, generating priority effects mediated by the host.

We tested whether the priming treatment altered the probability of a host becoming infected in the manipulated epidemic using a logistic mixed model. As predicted, hosts that were experimentally inoculated were more likely to become subsequently infected during the experimental epidemic (p = 0.0088; Fig. 2a; Table S1a). This effect was qualitatively similar using the (logit-transformed) proportion of leaves infected as a response measure representing infection severity (p = 0.019; Table S1b). Although host susceptibility to infection and the severity of infection were positively influenced by the priming treatment, this effect disappeared when we evaluated infection severity among infected hosts only (p = 0.82; Table S1c), suggesting that priority effects may act qualitatively (e.g., by altering susceptibility to infection) rather than quantitatively (e.g., by altering infection severity). This result is consistent with ecological theory, which suggests that priority effects should primarily function to prevent or facilitate establishment or persistence rather than population growth, *per se* (Fukami *et al.* 2016). This result therefore suggests that increased susceptibility to infection following early exposure to a pathogen strain can influence subsequent infection outcomes in the field.

**Figure 2.**
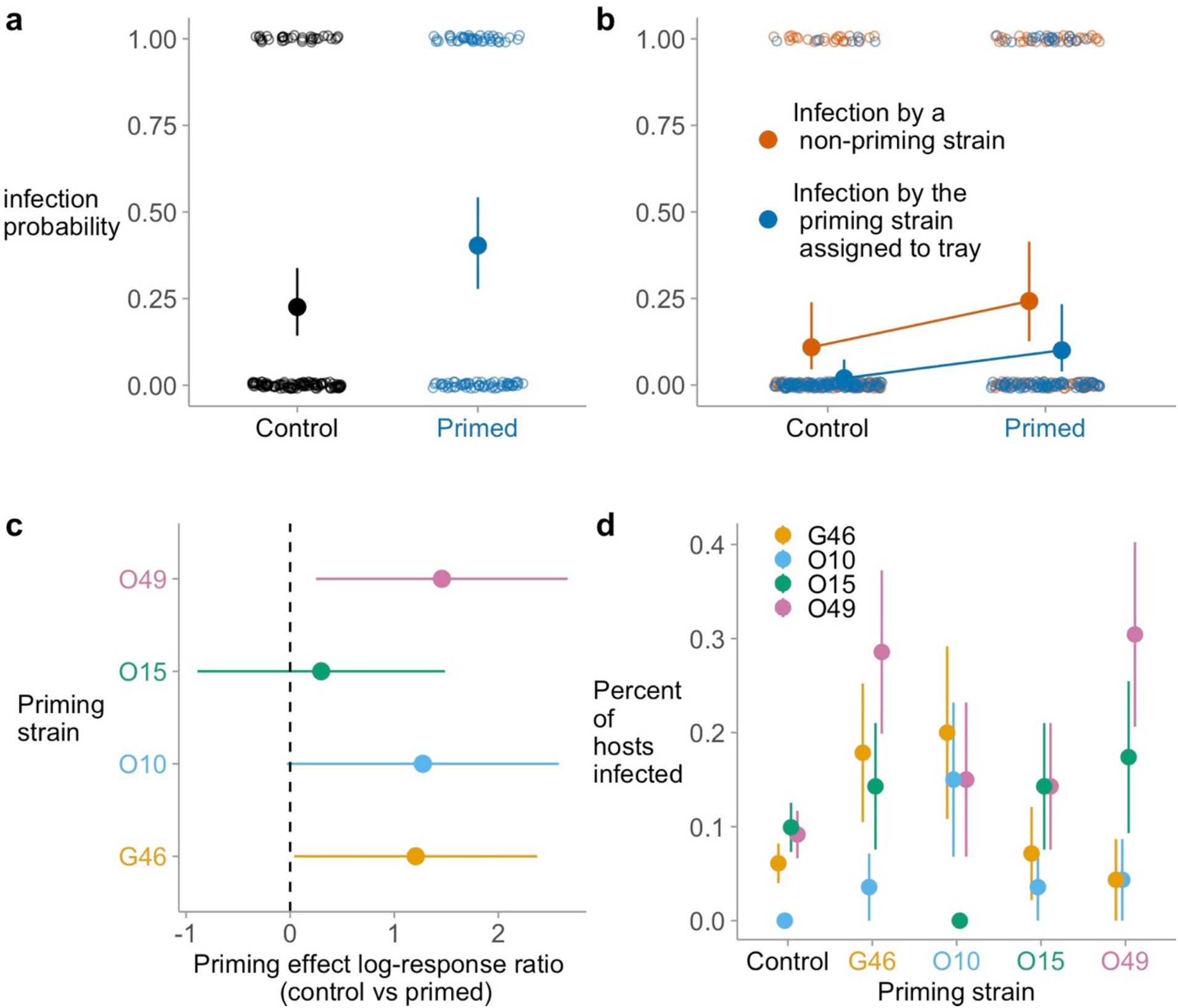
Results from the manipulated epidemic experiment. The effect of the priming treatment on a) whether or not a host became infected with any strain; b) whether or not a host became infected with the priming strain assigned to the experimental block (control hosts were mock-inoculated with the priming strain associated with that block), or any non-priming strain; c) whether or not a host became infected with any strain as a function of the priming strain, shown as a log-response ratio; d) the proportion of hosts infected by each strain during the manipulated epidemic experiment. In panels a-c, filled points are model-estimated means, error bars are model-estimated 95% confidence intervals, and open points show the raw data. In panel d, points show the proportion of hosts infected and the error bars show one standard error around those points. These results highlight the differential effect of each strain on the priming response of the plant as well as the differential sensitivity of each strain to that priming effect. Only strain O15 was a uniformly poor primer of subsequent infection. All other parasite strains significantly facilitated subsequent infection by at least one other strain. Together, these results suggest that by differentially determining the plant response to infection and experiencing differential sensitivity to that response, prior infection can strongly alter assembly of these pathogen strains.

We next tested whether the facilitative effect of early exposure on susceptibility to infection during the experimental epidemic differed among host genotypes and priming strains. Consistent with theory grounded in the history of interactions between host and parasite populations (e.g., Tollenaere *et al.* 2015), the facilitative effect of early infection depended on the priming strain (p = 0.050) and host plant genotype (p = 0.024), though there was no interaction between host plant genotype and the priming treatment (p = 0.86; Table S2a, Fig. 2c). We therefore dropped the non-significant interaction, resulting in a reduced model, and estimated the coefficients from the reduced model. Consistent with facilitative priority effects, the priming strains G46 and O49 significantly increased the probability of infection under field conditions (p = 0.039 and p = 0.016, respectively), while strain O10 marginally significantly increased the probability of infection (p = 0.051) and strain O15 did not (p = 0.62). This result suggests that only some early arrivers strongly influenced the availability of the niche for later arrivers, providing a possible mechanism for facilitative priority effects within hosts (e.g., Vannette & Fukami 2014). These results were qualitatively similar using the proportion of leaves infected as a metric of infection severity (Table S2b). Although the effect of the priming treatment on host susceptibility to infection and the severity of infection were influenced by host and parasite genotype, this effect disappeared when we evaluated infection severity among infected hosts only (Priming strain p = 0.95; Host plant genotype p = 0.61; Table S2c), lending further support to the idea that priority effects act qualitatively rather than quantitatively in this system.

Together, these results suggest that the host response to initial infection depended on which parasite strain arrived first. However, for priority effects to alter the structure of parasite assemblages, later arriving strains must also be sensitive to the plant response to initial infection (Vannette & Fukami 2014). To explore this mechanism of within-host interactions, we next genotyped infections on each individual host following exposure to all four strains under natural conditions and then tested for interactions among the priming treatment, plant genotype, and whether or not the later arriving strain was the same as the early arriving strain. For priority effects to occur, facilitative effects should occur among different strains. In other words, a priority effect could only occur if the early arriving strain facilitated other later arriving strains. Across both treatments, infection by a non-priming strain was about 1.4 times more likely than infection by the priming strain (p < 0.001; Fig 2b). Consistent with the hypothesis that parasite strains can exhibit within-host priority effects, the effect of early infection on the probability of subsequent infection was qualitatively similar between secondary infections caused by the priming strain (p = 0.004) and secondary infections caused by a different strain from the priming strain (p = 0.033). In other words, there was a significant main effect of the priming treatment (p = 0.003), but no interaction between the priming treatment and whether or not the host became infected with a strain other than the priming strain (p = 0.24; Table S3).

Finally, we tested whether within-host priority effects altered the structure of parasite assemblages using a multivariate generalized linear model (Wang *et al.* 2012; Warton *et al.* 2012). As expected, different parasite assemblages formed on hosts that received different priming treatments (LRT = 37; p = 0.040; Table S4; Fig. 2d), though there were no significant differences among different host genotypes (LRT = 34.94; p = 0.085), and the effect of priming treatments on the structure of parasite assemblages did not interact with host genotype (LRT = 57; p = 0.077). Thus, priority effects among strains altered parasite assembly, and the trajectory of assembly depended on the identity of the early arriving strain, but not the genotype of the host.

### Can host-mediated priority effects among parasites influence parasite assembly during a natural epidemic?

The manipulated epidemic experiment tested whether hosts could mediate priority effects among parasites, and whether such priority effects could influence parasite assembly during an experimental epidemic. We next carried out the natural epidemic experiment (Fig 1) to test whether host-mediated priority effects could be generalized to predict the outcome of natural epidemics by embedding sentinel hosts that were either primed or mock-inoculated into an ongoing epidemic in three wild host populations in the Åland archipelago. In addition to manipulating infection sequence (primed vs mock-inoculated), this experiment also manipulated the timing of prior infection by priming hosts either four or eight days prior to exposing hosts to the natural epidemic.

We first tested whether the host response to prior infection could generate priority effects using a logistic mixed model. Consistent with expectations from the manipulated epidemic experiment, the priming treatment significantly influenced the probability of a host becoming infected during the natural epidemic experiment (p < 0.001; Table S5). However, in contrast with the manipulated epidemic experiment, there was no significant effect of host genotype (p = 0.34). Consistent with expectations grounded in previous studies of this system (Penczykowski *et al.* 2018), the probability of infection differed among populations (p = 0.002). But, in contrast with expectations grounded in laboratory studies of plant immunity (Pieterse *et al.* 2012) and ecological theory (Fukami 2015), there was no difference in the magnitude of the priming effects between hosts that were primed eight days prior to experimental placement in the field and hosts that were primed four days prior to placement in the field (Fig 3a). The results were qualitatively similar using the (logit-transformed) proportion of leaves infected as a response variable representing infection severity. However, there was also a significant three-way interaction in the model of infection severity, suggesting that priority effects occurred, that these priority effects depended on infection sequence and timing, but only in certain populations, and only among certain host genotypes (Fig S1a). The reduced model of infection severity among infected hosts also included significant two-way interactions between population and host genotype (p = 0.033), and between host genotype and experimental treatment (p = 0.019; Fig S1b).

**Figure 3.**
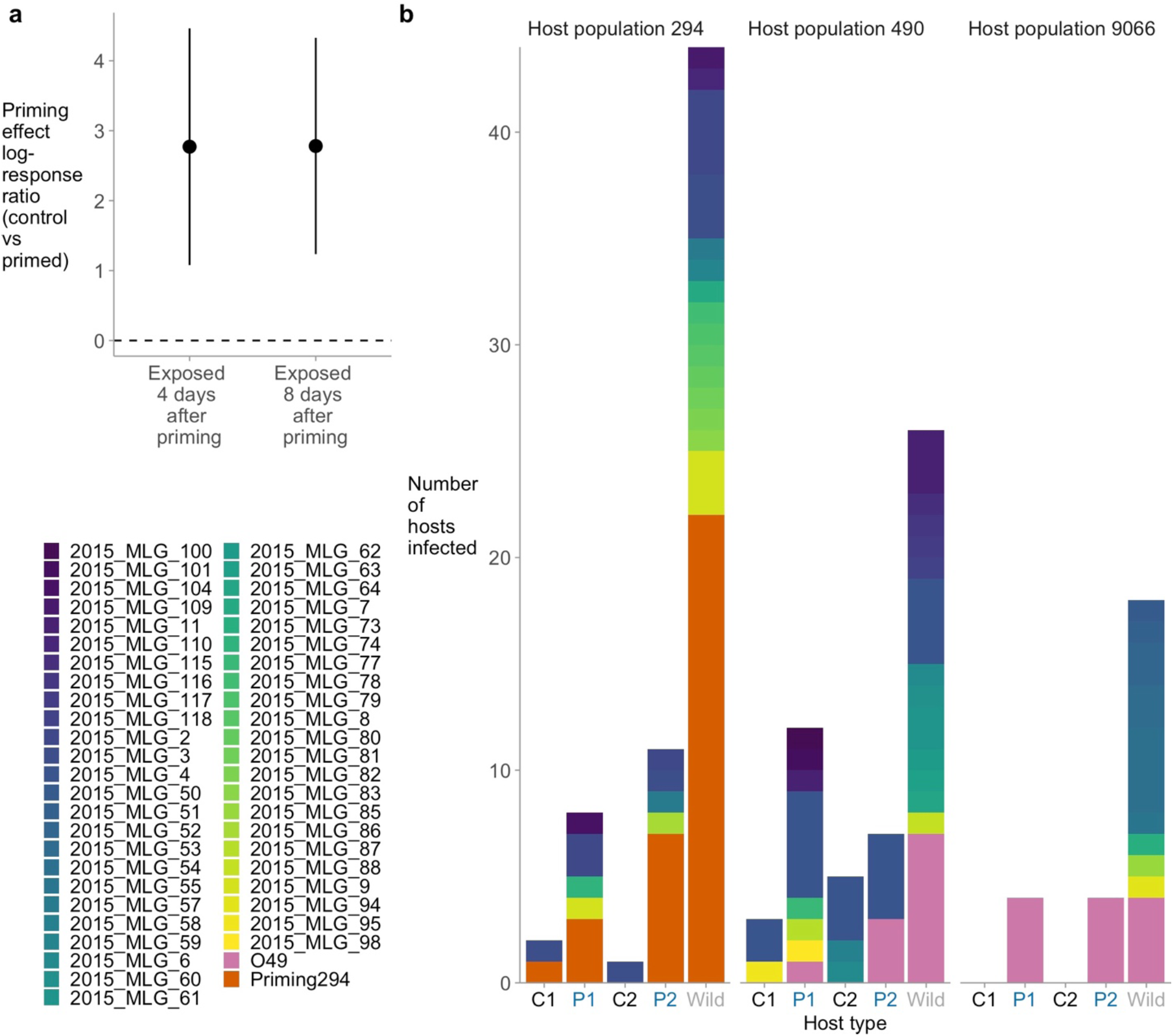
Results from the natural epidemic experiment. The effect of the sequence and timing of infection on a) whether or not a host became infected with any strain, shown as a log-response ratio between each control and each priming treatment. (e.g., C1 vs P1 and C2 vs P2); and b) the number of hosts infected by each strain grouped by whether a host was experimental (C1, C2, P2, P2) or wild (shown in grey). Plants were primed either 8 days (treatment P1) or 4 days (treatment P2) prior to being placed into the field. There were also C1 and C2 control plants set up at the same time (but mock inoculated). Points are model-estimated means, and error bars are model-estimated 95% confidence intervals. Colors represent different strains, with pink and orange corresponding to the two priming strains used in this experiment (O49 and Priming294, respectively). These results show that, consistent with the manipulated epidemic experiment, experimental priming increased the probability of a host becoming infected. However, that effect did not depend on infection timing (P1 vs P2). As a consequence of this effect, experimentally primed hosts had more complex parasite communities that were more similar to wild hosts than mock-inoculated control hosts.

Our result that host-mediated interactions among parasites almost universally favored coinfection is in contrast to prior studies suggesting that priming can commonly reduce the probability of coinfection through cross resistance (Fulton 1986; Pieterse *et al.* 2014; Biere & Goverse 2016), potentially raising concerns that these results might be system specific; however, a prior study in this wild plant pathosystem suggests a different explanation for these contrasting results. Specifically, using different *Plantago lanceolata* hosts and *Podosphaera plantaginis* genotypes, Laine (2011) found that priming reduced spore production in the lab, but increased infection severity in the field. Thus, we suggest that the difference between the results presented here and commonly reported results of cross resistance in other systems might be attributable to laboratory versus field environments, adding to a growing body of evidence that within-host interactions studied in the laboratory might be poor predictors of infection outcomes during natural epidemics (Seabloom *et al.* 2009; Leung *et al.* 2018; Clay *et al.* 2019).

Finally, we tested whether priority effects among parasite strains could lead to variation in the structure of parasite assemblages among hosts during a natural epidemic using a multivariate generalised linear model. This model did not include any significant interactions. However, as expected, there were different parasite assemblages on hosts that received different priming treatments (LRT = 111; p < 0.001), among different populations (LRT = 90; p < 0.001), and among different host genotypes (LRT = 44; p = 0.048; Table S6; Fig 3b). Thus, host genotype, spatial structure, and priority effects among strains all independently altered parasite assembly in the natural epidemic experiment.

### Can a signal of host-mediated facilitative priority effects among parasites be detected in natural populations?

Our experimental results showed consistent host-mediated facilitative priority effects among parasite strains. However, in addition to host-mediated interactions, parasites can also interact via resource or interference competition during natural epidemics (Mideo 2009). In theory, the host-mediated interactions can be either antagonistic or facilitative, whereas resource and interference competition are generally expected to reduce the risk of coinfection (Mideo *et al.* 2008; Pedersen & Greives 2008; Halliday *et al.* 2018). Thus, although both experiments suggested that the host response to prior infection can facilitate subsequent infection via within-host priority effects, the degree to which this process plays out to influence parasite assemblages during natural epidemics remains unclear.

We tested whether host-mediated priority effects are sufficiently important to influence the structure parasite assemblages in nature by analyzing the results of a longitudinal survey of wild host individuals during a natural epidemic (i.e., the wild host survey; Fig 1). The wild host survey was carried out in the Åland archipelago, and included 105 host individuals from 13 populations, sampled biweekly for infection starting on 7 July, 2014. Once a host became infected, it entered the dataset as a focal host. To determine infection sequence among hosts, we sampled lesions and genotyped infections twice on each focal host: first when more than one leaf on a focal host was infected, and then again at the end of the season. These two genotyping sessions provide data on the sequence and timing of infection among hosts, while biweekly surveys of whole host populations provide information on parasite phenology.

We hypothesized that if the host response to prior infection was sufficiently strong to alter parasite community assembly, then we would observe a signal of facilitation among co-occurring parasite strains during a natural epidemic. To test whether parasite strains exhibit priority effects within hosts, we first fit a series of cox proportional hazards models following Halliday et al (2017, 2018). These models test whether the time until infection by each parasite strain was influenced by whether or not a host had been previously infected by another strain. In total this analysis tested for 286 potential pairwise interactions among parasite strains. However, despite the large number of potentially interacting parasite strains, only three pairwise interactions resulted in a significant priority effect as defined by Halliday et al (2017, 2018). Specifically, 2014_MLG_78 and 2014_MLG_10 were significantly facilitated by prior infection by 2014_MLG_5 and 2014_MLG_42 respectively (p = 0.009; p = 0.035), while 2014_MLG_3 was antagonized by prior infection by 2014_MLG_43 (p=0.009); importantly, given the numerous comparisons per host individual, even these significant effects should be interpreted with caution. These results therefore suggest that, on average, parasite strains were not exhibiting measurable priority effects within hosts.

There are many potential reasons for the absence of significant interactions among parasites, including the physiological state of wild host plants (Penczykowski *et al.* 2018), the presence of counterbalancing direct interactions among parasites (Mideo *et al.* 2008), skewed distribution of infections by the different strains in the field (Laine 2007), or the relatively infrequent sampling of host individuals in the field. However, although within-host priority effects were rarely significantly positive or negative, we still wanted to know whether a signal of facilitation could be observed among potentially interacting parasite strains. Overall, early infection tended to facilitate subsequent infection by other strains more commonly than preventing subsequent infection by other strains (p<0.001; Fig 4a), consistent with priority effects being mediated by the host response to prior infection. These results support the hypothesis that facilitative interactions among parasite strains, mediated by the host response to prior infection, would result in a signal of facilitation among co-occurring parasite strains.

**Figure 4.**
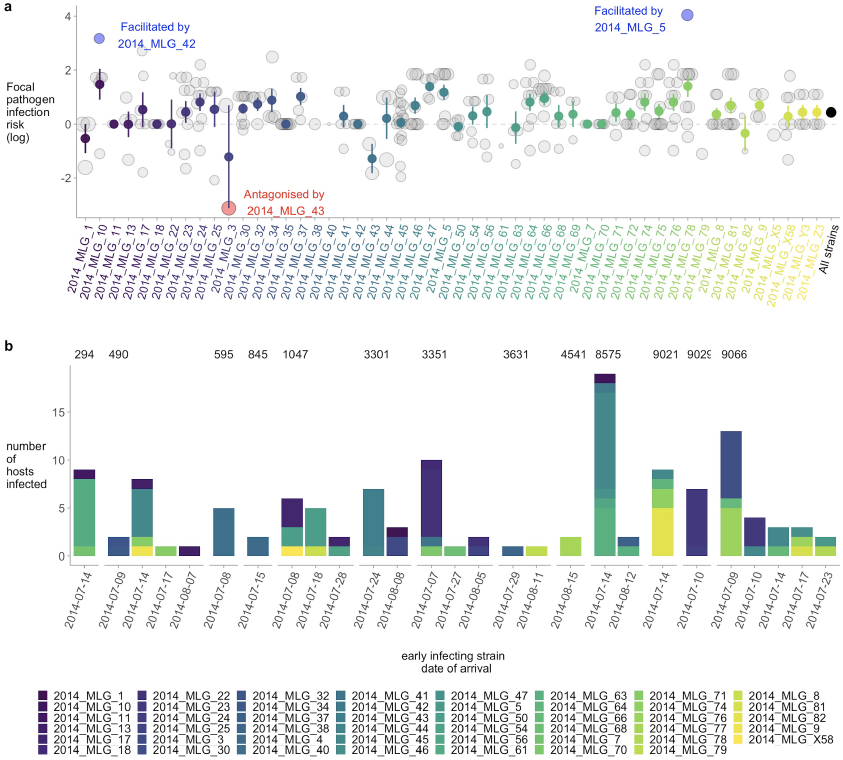
Results from the wild host survey. a) The effect of infection sequence on the risk of a host becoming infected with each parasite strain. The x-axis shows the focal (i.e., late arriving) parasite strains from field-collected samples. The grey, blue, and red points are coefficient estimates from cox proportional hazards models measuring the pairwise interaction between each focal strain and each co-occurring other (i.e., early arriving) strain; point size represents the average number of surveys per host individual; blue and red indicate significant facilitation and antagonism (p < 0.05), respectively, and grey indicates insufficient evidence for significant priority effect (p > 0.05). The colored points with error bars show the mean and one standard error across all coefficients for each focal parasite strain. The black point is the model estimated mean across all parasite strains, and the black error bar (which is small and largely obscured by the black circle) is the 95% confidence interval from an intercept-only model, indicating that on average, interactions tended to be positive. b) The number of hosts infected by each strain at the end of the growing season grouped by the date that the early arriving strain was first observed in the field. Panels represent different host populations. These results show that, across all parasite strains, interactions were most commonly facilitative, and hosts that were first infected by strains with early phenology were more likely to become infected by more complex communities at the conclusion of the growing season.

Finally, to test whether parasite phenology among strains altered parasite assembly within hosts, we fit a multivariate generalised linear model. We hypothesized, in accordance with ecological theory (e.g., Vannette & Fukami 2014) and our experimental results, that strains that emerged later in the growing season (i.e., strains with later phenology) would be more sensitive to facilitative priority effects, and that strains that emerged earlier in the growing season (i.e., strains with early phenology) would more strongly influence parasite assembly. Consistent with this hypothesis, the structure of parasite assemblages differed significantly among hosts with differing phenology of the early-arriving strains (LRT = 15.23; p = 0.002; Fig 4b; Table S7), even after accounting for survey date (LRT = 238; p = 0.001), and host different population (LRT = 189; p = 0.001).

## Conclusions

The sequence and timing of infection can strongly influence parasite interactions and epidemics (Halliday *et al.* 2017, 2018; Clay *et al.* 2019, 2020), yet the degree to which this process is driven by the host response to infection versus other mechanisms of interaction among parasites remains poorly understood. This study leveraged a model wild-plant pathosystem to fill this gap (Penczykowski *et al.* 2016). Specifically, our study revealed three key findings: (1) by manipulating infection sequence during an experimental epidemic, we showed that host-mediated interactions among parasites almost universally favored coinfection; (2) by manipulating infection sequence during a natural epidemic, we showed that this process could alter how parasite strains assemble; and (3) by tracking wild host individuals during the course of a different natural epidemic, we identified a signal of host-mediated facilitation among parasite strains that could be linked to the structure of parasite assemblages. Our results therefore provide comprehensive evidence that parasite interactions, mediated by the host response to initial infection, can facilitate subsequent infection by different parasite strains, altering the trajectory of parasite assembly during natural epidemics.

In addition to isolating host-mediated interactions from other interactions among parasites, our experimental approach overcomes an additional key limitation of past observational studies: the need to rely on counterfactual reasoning. Linking infection sequence and within-host priority effects during natural epidemics is notoriously challenging (Budischak *et al.* 2018; Rynkiewicz *et al.* 2019). Our experimental data showed clear evidence of host-mediated facilitation among parasites. Pairing this experimental approach with a fine-scale survey and sampling of infections in the wild during a natural epidemic thus allowed us to interpret a signal of host-mediated facilitation that would be uninterpretable from observational data alone. Our results suggest that, by differentially determining the plant response to infection and experiencing differential sensitivity to that response, prior infection can strongly alter the structure of parasite assemblages during epidemics.

Priority effects among strains favored coinfection and altered parasite assembly, and the trajectory of assembly depended on the genotype of the early arriving strain. This result supports the idea that species interactions – in this case host-parasite and parasite-parasite interactions – can depend on intraspecific variation in characteristics of organisms (Bolnick et al. 2011; Laughlin et al. 2012; Siefert 2012). Intraspecific diversity is ubiquitous in host and parasite populations, and has prompted considerable research into local adaptation among hosts and parasites (Greischar & Koskella 2007), parasite aggregation (Shaw & Dobson 1995), and disease emergence (Lloyd-Smith et al. 2005). Yet, how this diversity impacts parasite assembly is not known. Importantly, our results suggest that by influencing host susceptibility, intraspecific variation can determine how strain diversity is spatially and temporally distributed during epidemics.

In our experimental manipulation of a natural epidemic, we found evidence that infection sequence, but not infection timing altered future host susceptibility. We also found strong evidence that host genotype and spatial structure altered the structure of parasite assemblages within hosts, but these effects did not alter the direction or magnitude of priority effects. Thus, we conclude that differences among host genotypes and among populations may play a large role in the assembly of parasite strains in natural populations, but these effects are independent of the robust effect of facilitation by sequentially arriving parasite strains in this system.

## Methods

### Study system

The host plant, *Plantago lanceolata* L. (ribwort plantain), is a perennial rosette-forming herb that reproduces either sexually, as an obligate outcrosser with wind-dispersed pollen, or via vegetative propagation of side-rosettes (Sagar & Harper 1964; Ross 1973). This species has a cosmopolitan distribution, and grows in fragmented populations on dry meadows and pastures in the Åland archipelago (SW Finland) (Ojanen *et al.* 2013). Plantago lanceolata is host to the obligate parasite *Podosphaera plantaginis* (Castagne; U. Braun and S. Takamatsu), a powdery mildew fungus in the order Erysiphales within the Ascomycota. The fungus grows on the leaf surface and extracts plant nutrients via haustoria that enter the epidermis (Bushnell 2002). On the leaf surface, mycelia produce chains of asexual, wind-dispersed transmission spores (conidia). The parasite reproduces clonally throughout the summer growing season, and then produces resting structures (chasmothecia) via haploid selfing or outcrossing (Tollenaere & Laine 2013) which enable the parasite to survive when the host plant has died back to rootstock in winter. Within the chasmothecia, haploid ascospores develop, which re-initiate epidemics in spring (Tack & Laine 2014).

The metapopulation dynamics of this host-parasite interaction in ca. 4000 populations in Åland have been studied since the year 2001 (Laine & Hanski 2006; Jousimo *et al.* 2014). Powdery mildew infection combined with stressful environmental conditions can cause high mortality of *Pl. lanceolata* (Laine 2004), and infection reduces host population growth rates (Laine 2004; Penczykowski *et al.* 2015).

Successful infection is the outcome of a high degree of specificity where a given *Pl. lanceolata* genotype can be susceptible to some *Po. plantaginis* strains while able to block infection by others (Laine 2004, 2007). Hosts that are qualitatively susceptible to a given parasite strain may still vary in their ability to mitigate its sporulation once infected (i.e., quantitative resistance). Evidence for diversity within and among host populations comes from laboratory inoculation experiments showing variation in resistance to a given set of parasite strains among clonal plant lines under controlled conditions (Laine 2004, 2007). The *Po. plantaginis* populations in Åland are also diverse, comprised of genetically and phenotypically distinct parasite strains (Tollenaere *et al.* 2012), with a high proportion of coinfection by different multilocus genotypes (MLGs) (Susi *et al.* 2015).

### Study design

The study consisted of two experiments and a fine-scale survey and sampling of infections in the wild (Fig. 1). In the first experiment, which we refer to as the “manipulated epidemic experiment”, we tested whether parasite strains exhibit priority effects that are mediated by the host plant response to prior infection. We addressed this question by first inoculating or mock-inoculating hosts of four different genotypes with one of four parasite strains and then later exposing the same hosts to all four strains in a common garden environment. In the second experiment, which we refer to as the “natural epidemic experiment”, we tested whether host-plant mediated priority effects among parasite strains can influence parasite assembly during a natural epidemic. We addressed this question by first inoculating or mock-inoculating hosts with one of three different parasite strains, then embedding those hosts into one of three wild host populations in order to expose the hosts to a natural epidemic, and then compared the effect of these treatments on the structure of the resulting parasite assemblage on each sentinel host plant as well as wild plants from the same populations. These experiments revealed strong facilitative priority effects among parasite strains, and indicated that these facilitative priority effects can alter parasite assembly during natural epidemics. Finally, we analyzed the results of an observational study, which we refer to as the “wild host survey,” to test whether a signal of host-plant mediated facilitative priority effects is detectable among parasite strains in natural populations. We addressed this question by tracking infections on wild host plants over the course of a natural epidemic.

#### Experimental plant genotypes

To test whether the outcome of sequential infections varied across different host plant genetic backgrounds, we used a set of four host plant maternal lines in both experiments (Table S8). Each maternal line came from a single seed head from a different mother plant in the Åland archipelago. Seeds were sown in 9 × 9 cm flower pots in a mixture of 30% sand, and 70% potting soil. Plants were maintained in the greenhouse at +20 °C until transport to the common garden location, where they were acclimated to outside conditions for at least two weeks before the start of the experiments.

#### Experimental parasite strains

The parasite strains used for inoculating plants in both experiments were collected from field populations in the Åland archipelago at the end of epidemics in 2014 (Table S9). We collected and purified the powdery mildew isolates as follows. Infected leaves were detached using forceps and placed into 9 cm Petri dishes containing moist filter paper. Between every sampled leaf, the forceps were sanitized with DNA-Away (Molecular Bio Products) to avoid cross-contamination. To ensure that each parasite isolate was a single strain (multi-locus genotype, MLG), we purified the isolates through three successive single-colony transfers of spores onto detached, greenhouse-grown leaves (Nicot *et al.* 2002).

Inoculated leaves were maintained on moist filter paper in Petri dishes in a growth chamber under standard conditions of 21°C (± 2 °C) and 16L:8D photoperiod. We then amplified the fungal isolates through 2-3 rounds of inoculations to generate enough spores for the experiments described below.

#### Manipulated epidemic experiment

We performed the manipulated epidemic experiment to test how inoculating a single leaf of the rosette with a single parasite strain (“priming”) affected susceptibility of the plant to later-arriving parasite strains. The common garden experiment was performed in a 30 × 45 m fenced field at the University of Helsinki’s Lammi Biological Station (Lammi, Finland). The experiment consisted of a total of 320 plants placed in groups of eight in plastic trays (0.5 × 0.3 m), with two plants from each of the four maternal plant lines in each of 40 trays. Trays were equally spaced apart in the field along a 5 m x 5 m grid. Each plant included two leaves from the same rosette spiral enclosed in separate sleeves made from spore-proof polyester material (pollination bags from PBS International) and secured at the leaf base, which were used for a separate study and are not discussed further. One plant from each maternal line in the tray was assigned to the “primed” treatment, and the other plants were assigned to the mock-inoculated “control” treatment. On 26 July 2015, primed plants were enclosed in a plastic bag with a single leaf emerging through a small hole in the bag. A fine paintbrush was used to inoculate that leaf with one of the four priming strains, depending on the tray ID. Twenty-four hours after inoculation, the leaf was covered with a spore-proof sleeve, the plastic bag was wiped with ethanol and removed. Control plants underwent the same procedure, but no powdery mildew was inoculated.

The plants in the common garden were then bulk-exposed to all four powdery mildew strains over the course of four days (30 July-2 August; days 4 to 8 post-priming). This was done by rotating heavily infected source plants next to the trays, such that each tray was exposed to each of the strains for 24 hours.

The plants in the common garden were screened for infection between 19-23 August. We recorded the total number of uninfected and infected leaves for each plant. We collected several infected leaves from each plant for genotyping. These infected leaves were stored in paper envelopes at room temperature until DNA extraction (see Genetic analyses section below).

#### Natural epidemic experiment

This experiment tested whether the sequence and timing of priming influenced subsequent infection among sentinel plants placed into three field populations during natural epidemics. We used the same set of four host plant maternal lines as in the common garden experiment, and plants in the priming treatment were inoculated with a parasite strain that had been present in the field population the previous year (Table S8). Priming of a single leaf was performed as in the common garden experiment. To manipulate infection timing, plants were primed either 8 days (treatment P1) or 4 days (treatment P2) prior to being placed into the three field populations on 4 August 2015. There were also C1 and C2 control plants set up at the same time (but mock inoculated). For each of the four host plant maternal lines in each of the three field populations, we had 10 replicates of P1, 10 replicates of P2, 5 replicates of C1, and 5 replicates of C2, for a total of 360 plants. Groups of paired primed and control sentinel plants were placed on plastic trays throughout the field populations. The trays were watered and moved to new locations every two days for 8 days, to standardize exposure to powdery mildew spores. After the 8 days of exposure to natural epidemics, the plants were covered individual spore-proof pollination bags and transported back to the Lammi Biological Station, where infections were allowed to continue developing for another 10 days. Then we counted infected and uninfected leaves in each size class as described for the common garden experiment. We also saved infected leaves in individual paper envelopes to genotype and determine which strains infected them.

While the sentinels were in the field populations, we surveyed wild plants from each population for infection and tagged infected plants located at least 1.5 m apart. Up to 44 infected plants per population were tagged. At the end of the epidemics, we collected samples of infected leaves from the tagged plants for genotyping and parasite strain identification. Infected leaves were stored in paper envelopes at room temperature until prepared as samples.

#### Wild host survey

The wild host survey was carried out in the Åland archipelago in 2014 and consisted of fifteen host populations that had been infected for at least three consecutive years (Jousimo *et al.* 2014). Distances between pairs of populations ranged from ∼ 1 km to ∼ 40 km. Each population was surveyed biweekly for infection starting on 7 July, 2014, by visually scanning plants for signs of the mildew. When host individuals became infected, the date of infection was recorded and those hosts were physically tagged in order to be resurveyed as focal hosts. Up to thirty focal hosts were tagged in each population. Focal plants were located at least 3 m from one another and their locations were recorded by GPS. To minimize the impact of sampling on pathogen community assembly, we sampled lesions from focal hosts and genotyped the infections only after more than one leaf of a focal host was infected. We sampled lesions in such a manner that spores also remained on the plants, thus the infection was not removed from the epidemic. All hosts that survived were then resampled at the end of the season (n = 105 hosts across 13 populations). Sampling consisted of placing infected leaves in paper envelopes, which were stored in a cool, dry place until the end of the field season, at which point the samples were taken to Helsinki and stored at -20C. These two genotyping sessions were used to infer the sequence of infection on host individuals, while the frequent surveys of whole host populations were used to infer phenology of the parasite strains.

#### Genetic analyses

We genotyped infections in all three studies to determine which powdery mildew strains successfully established on the host plants. Each sample consisted of a lesion from an infected leaf, which we placed into a 1.5 mL tube that was stored at -20°C until DNA was extracted using an E.Z.N.A. Plant Mini Kit (Omega Bio-Tek, Norcross, GA) at the Institute of Biotechnology, University of Helsinki. The lesions consisted of both host tissue and fungal material. Samples were genotyped at 19 single nucleotide polymorphism (SNP) loci with the Sequenome iPlex platform at the Institute for Molecular Medicine Finland (See Tollenaere *et al.* 2012; Parratt *et al.* 2017 for details). Because *Po. plantaginis* conidial spores are haploid, samples were classified as coinfected if two different nucleotides were called at any locus (Tollenaere *et al.* 2012). The observed coinfections were resolved into single infections with an algorithm that compared each coinfection profile to the genotypes of all single infections in the experiment (i.e., to the four strains in the manipulated epidemic experiment or to all single infections from the same population in the natural epidemic experiment). When a match was found, the genotype of the other coinfecting strain could be determined as having the complementary alleles at the heterozygous loci. However, for samples with only a few heterozygous loci and where multiple strains had the same nucleotides at those loci, we could only unambiguously identify one of the two coinfecting strains. For samples from the manipulated epidemic experiment that failed to call all 19 SNPs, we were still able to identify the strain if the nucleotides at the successfully called SNPs were unique to one of the four strains in that experiment. However, samples from the natural epidemic experiment or wild host surveys that were missing genotype data from any of the 19 SNPs were excluded from the analysis.

From sentinel plants in the manipulated epidemic and natural epidemic experiments, we randomly selected four infected leaves per plant for genotyping (if fewer than four leaves on the plant were infected, then all infected leaves were sampled). In addition, we genotyped infections from a subset of the primed leaves to verify that plants were primed with the correct parasite strains. A large number of samples failed to call several of the 19 SNP loci during our first round of genotyping in spring 2016. To replace those samples for which genotyping failed, we extracted DNA from remaining infected leaves from the same plants and genotyped those replacement samples in spring 2017.

### Analysis

All analyses were conducted in R version 3.5.2 (R Core Team 2015). We omitted plants from analyses that were inoculated, but never became infected or that were mock-inoculated but became infected (85 in the manipulated epidemic experiment; 165 in the natural epidemic experiment), as well as plants that died prior to data collection (4 in the manipulated epidemic experiment; 2 in the natural epidemic experiment) resulting in a total sample size of n = 231 in the manipulated epidemic experiment and n = 193 in the natural epidemic experiment.

#### Manipulated epidemic experiment

We first tested whether the priming treatment altered infection during the manipulated epidemic by constructing three models using the R package lme4 (Bates *et al.* 2014): (1) the probability of a plant becoming infected, using a logistic mixed model, (2) the logit-transformed proportion of leaves infected as a response measure representing infection severity, using a linear mixed model, and (3) the logit-transformed proportion of leaves infected as a response measure representing infection severity, limited to infected hosts only, using a linear mixed model. All three models included the experimental treatment (inoculated vs. mock inoculated) as a fixed effect. We included the log-transformed number of leaves as a fixed covariate in the model, because plants with more leaves have a higher probability of intercepting infectious spores. Inoculation tray was included as a random effect in the model. To test whether treatment effects differed among host genotypes and priming strains, we fit three models with the same three response variables, this time including the full priming treatment (five levels: mock-inoculated, and four priming strains), plant genotype, and their interactions as fixed effects, the log-transformed total number of leaves as a fixed covariate, and priming tray as a random effect. We evaluated differences among various model coefficients using the emmeans package in R (Lenth *et al.* 2018).

To test whether the priming strain facilitated other later arriving strains at least as strongly as it facilitated itself, we fit a model with a binomial response representing whether or not the host became infected, and including the priming treatment as a binary factor (control vs primed) and its interaction with whether or not the infecting strain was different from the priming strain. The model also included log-transformed total number of leaves as a fixed covariate and the tray and plant id as nested random intercepts. Finally, we tested whether within-host priority effects altered the structure of parasite assemblages, using a multivariate generalized linear model using the MVabun package in R (Wang *et al.* 2012). To measure the effect of prior infection on the structure parasite assemblages, we constructed a model that tests whether the distribution of infection by each strain was affected by the priming treatment (five levels: control, and four priming genotypes), plant genotype, and their interactions, including the log-transformed total number of leaves as a covariate. These models avoid some of the problems associated with distance-based models such as permANOVA. However, because these models cannot handle unbalanced grouping variables, experimental tray was included as a covariate rather than a random effect in the model.

#### Natural epidemic experiment

We first tested whether the priming treatment altered infection during the natural epidemic by constructing three models using the lme4 package in R: (1) the probability of a plant becoming infected, using a logistic mixed model, (2) the logit-transformed proportion of leaves infected as a response measure representing infection severity, using a linear mixed model, and (3) the logit-transformed proportion of leaves infected as a response measure representing infection severity, limited to infected hosts only, using a linear mixed model. Each model included host population, plant genotype, and the experimental treatment (C1: control for the first priming treatment, C2: control for the second priming treatment, P1: first priming treatment at eight days prior to placement in the field, and P2: second priming treatment at four days prior to placement in the field) as interactive fixed effects and the log-transformed total number of leaves as a fixed covariate. Experimental tray was included as a random intercept in the model, but this random effect did not explain any of the variance in some of the models, leading to a computational singularity. Despite the computational singularity, we opted to keep experimental tray in all models to account for non-independence among samples in each patch. In the logistic regression model, there were no significant interactions (p = 1), and owing to the binary response variable and high number of predictors, the model suffered from complete separation. We therefore iteratively removed all non-significant interactions, yielding a reduced model. We evaluated differences among various model coefficients in the reduced models using the emmeans package.

We tested whether priority effects among parasite strains could lead to variation in the structure of parasite assemblages among hosts during a natural epidemic with a multivariate generalised linear model using the MVabun package in R, to test whether the distribution of infection by each strain was affected by the priming treatment (four levels: C1, C2,P1, and 2), patch, plant genotype, and their interactions, including the total number of leaves as a covariate. To fit this model, we removed from the dataset any plant that had an infection that could not be genotyped, resulting in a total of 181 plants for this analysis. Similar to previous analyses, we reduced the model by removing non-significant interactions.

#### Wild host survey

To test whether parasite strains exhibit priority effects within hosts, we first fit a series of cox proportional hazards models following Halliday et al (2017, 2018) using the coxphf package in R (Ploner & Heinze 2015). To fit these models, we made one critical assumption about the data: we assumed that whichever strain was first observed in genotyping was the “early arriving strain”. This assumption allows us to use temporal data from the first observations of a host, regardless of the time between that survey and the genotyping date. However, this assumption ignores the possibility of rare parasite strains being locally cleared early during the epidemic. Furthermore, because we could not resolve the sequence of infection on host individuals that are coinfected during the initial genotyping survey (occurring in 22/105 host individuals), both coinfecting parasites were assumed to have been present at the initial infection of the host. We then fit a series of cox proportional hazards models of infection by each parasite as a focal (i.e., late-arriving) parasite with prior infection by each other (i.e., early-arriving) parasite as the only predictor in each model. These models tested whether the time until infection by each strain was influenced by infection sequence on a host individual. Across the Åland archipelago, distinct host populations often harbor distinct parasite assemblages. Parasite strains were therefore only modeled among hosts that occurred in populations where those Parasites had been observed. To explore whether the magnitude of priority effects among parasite strains tended to be facilitative or antagonistic, we next fit an intercept only linear mixed model with the coefficient from the cox proportional hazards models (i.e., the interaction coefficient) as the response variable and the identity of the focal (i.e., late arriving) parasite in the cox proportional hazards models as a random intercept, weighting the regression by the number of surveys per host individual to give more explanatory power to host individuals that were surveyed more times over the growing season. Finally, we tested whether parasite phenology among strains altered the structure of parasite assemblages within hosts using a multivariate generalised linear model on the distribution of infections by each strain at the end of the epidemic, in the R package MVabund. To measure parasite phenology, we recorded the earliest date that each strain was observed in the field during the 2014 epidemic. We then modeled the presence or absence of each strain at the end of the season as a function of the phenology of the early-infecting strains, with the sampling date of the final survey and host population as covariates in the model.

