## Supplement for "Facilitative priority effects drive parasite assembly under coinfection"

**Supplemental materials**

**Table S1.** Analysis of Deviance (Type II Wald $\chi^2$ tests) testing the effect of the priming treatment on mildew infection.

| | Chisq | Df | p-value |
|---|---|---|---|
| *A) Probability of a host becoming infected* | | | |
| Total number of leaves (log) | 0.33 | 1 | 0.56 |
| Treatment (primed vs control) | 6.87 | 1 | 0.0088 |
| *B) Logit-transformed proportion of leaves infected* | | | |
| Total number of leaves (log) | 0.0015 | 1 | 0.97 |
| Treatment (primed vs control) | 5.46 | 1 | 0.019 |
| *C) Logit-transformed proportion of leaves infected among infected hosts only* | | | |
| Total number of leaves (log) | 1.9 | 1 | 0.17 |
| Treatment (primed vs control) | 0.051 | 1 | 0.82 |

**Table S2.** Analysis of Deviance (Type II Wald $\chi^2$ tests) testing whether the effect of the priming treatment on mildew infection differed among host genotypes and priming strains.

| | Chisq | Df | p-value |
|---|---|---|---|
| *A) Probability of a host becoming infected* | | | |
| Total number of leaves (log) | 0.33 | 1 | 0.56 |
| Priming strain (control, G46, O10, O15, O49) | 9.51 | 4 | 0.050 |
| Host genotype (511_14, 4_14, 511_18, 9031_19) | 9.43 | 3 | 0.024 |
| Priming strain × Host genotype | 6.95 | 12 | 0.86 |
| *B) Logit-transformed proportion of leaves infected* | | | |
| Total number of leaves (log) | 1.05 | 1 | 0.31 |
| Priming strain (control, G46, O10, O15, O49) | 9.33 | 4 | 0.053 |
| Host genotype (511_14, 4_14, 511_18, 9031_19) | 8.32 | 3 | 0.040 |
| Priming strain × Host genotype | 5.96 | 12 | 0.92 |
| *C) Logit-transformed proportion of leaves infected among infected hosts only* | | | |
| Total number of leaves (log) | 0.90 | 1 | 0.34 |
| Priming strain (control, G46, O10, O15, O49) | 0.72 | 4 | 0.95 |
| Host genotype (511_14, 4_14, 511_18, 9031_19) | 1.83 | 3 | 0.61 |
| Priming strain × Host genotype | 6.93 | 12 | 0.86 |

**Table S3.** Analysis of Deviance (Type II Wald $\chi^2$ tests) testing whether effect of early infection on the probability of subsequent infection was qualitatively similar between secondary infections caused by the priming strain and secondary infections caused by a different strain from the priming strain.

| | Chisq | Df | p-value |
|---|---|---|---|
| Total number of leaves (log) | 0.43 | 1 | 0.51 |
| Treatment (primed vs. control) | 8.99 | 1 | 0.003 |
| Secondary infection type (priming strain vs. non-priming strain) | 14.62 | 1 | 0.0001 |
| Treatment × Infection type | 1.36 | 1 | 0.24 |

**Table S4.** Analysis of Deviance calculated using 999 resampling iterations via parametric resampling (to account for correlation in testing) testing whether within-host priority effects altered the structure of parasite assemblages in the manipulated epidemic experiment using a multivariate generalized linear model.

| | Residual DF | DF difference | Deviance | p-value |
|---|---|---|---|---|
| Experimental tray (i.e., block) | 190 | 39 | 119.64 | 0.66 |
| Total number of leaves (log) | 189 | 1 | 4.38 | 0.50 |
| Priming strain (control, G46, O10, O15, O49) | 185 | 4 | 36.99 | 0.040 |
| Host genotype (511_14, 4_14, 511_18, 9031_19) | 182 | 3 | 34.94 | 0.085 |
| Priming strain × Host genotype | 170 | 12 | 56.78 | 0.077 |

**Table S5.** Analysis of Deviance (Type II Wald $\chi^2$ tests) testing whether the host response to prior infection could generate priority effects in the natural epidemic experiment, using a logistic mixed model. Higher-level interactions were omitted from this model as they lead to complete separation in the fixed effects

| | Chisq | Df | p-value |
|---|---|---|---|
| Total number of leaves (log) | 1.55 | 1 | 0.21 |
| Host population | 12.23 | 2 | 0.0022 |
| Host genotype | 3.32 | 3 | 0.34 |
| Treatment (primed vs. control) | 21.30 | 3 | <0.0001 |

**Table S6.** Analysis of Deviance calculated using 999 resampling iterations via parametric resampling (to account for correlation in testing) testing whether within-host priority effects altered the structure of parasite assemblages in the natural epidemic experiment using a multivariate generalized linear model.

| | Residual DF | DF difference | Deviance | p-value |
|---|---|---|---|---|
| Total number of leaves (log) | 179 | 1 | 16.77 | 0.181 |
| Host population | 177 | 2 | 89.56 | 0.001 |
| Treatment (C1, C2, P1, P2) | 174 | 3 | 110.71 | 0.001 |
| Host genotype (511_14, 4_14, 511_18, 9031_19) | 171 | 3 | 44.48 | 0.048 |
| Population × Treatment | 165 | 6 | 0.89 | 0.433 |
| Population × Host genotype | 159 | 6 | 5.84 | 0.178 |
| Treatment × Host genotype | 150 | 9 | 20.26 | 0.157 |
| Population × Treatment × Host Genotype | 134 | 17 | 0.02 | 0.574 |

**Table S7.** Analysis of Deviance calculated using 999 resampling iterations via parametric resampling (to account for correlation in testing) testing whether within-host priority effects altered the structure of parasite assemblages in the wild host survey using a multivariate generalized linear model.

| | Residual DF | DF difference | Deviance | p-value |
|---|---|---|---|---|
| Survey date | 96 | 8 | 238.26 | 0.001 |
| Host population | 86 | 10 | 188.78 | 0.001 |
| Phenology of the early-arriving strain | 91 | 1 | 15.23 | 0.002 |

**Table S8.** Experimental plant genotypes

| Plant line name | Population of origin of mother plant |
|---|---|
| 4_14 | 4 |
| 511_14 | 511 |
| 511_18 | 511 |
| 9031_19 | 9031 |

**Table S9.** Experimental parasite strains

| Powdery mildew strain name | Population of origin | Year strain isolated | Common garden trays primed with this strain (tray IDs) | Sentinel groups primed with this strain (population names) |
|---|---|---|---|---|
| O10 | 9029 | 2014 | 1, 3, 5, 9, 14, 15, 20, 22, 30, 35 | -- |
| O15 | 3301 | 2014 | 4, 8, 12, 24, 25, 29, 31, 33, 34, 36 | -- |
| G46 | 475 | 2014 | 2, 7, 11, 16, 17, 18, 28, 32, 38, 40 | -- |
| O49 | 490 (also present in 9066) | 2014 | 6, 10, 13, 19, 21, 23, 26, 27, 37, 39 | 490, 9066 |
| Priming294 | 294 | 2014 | -- | 294 |

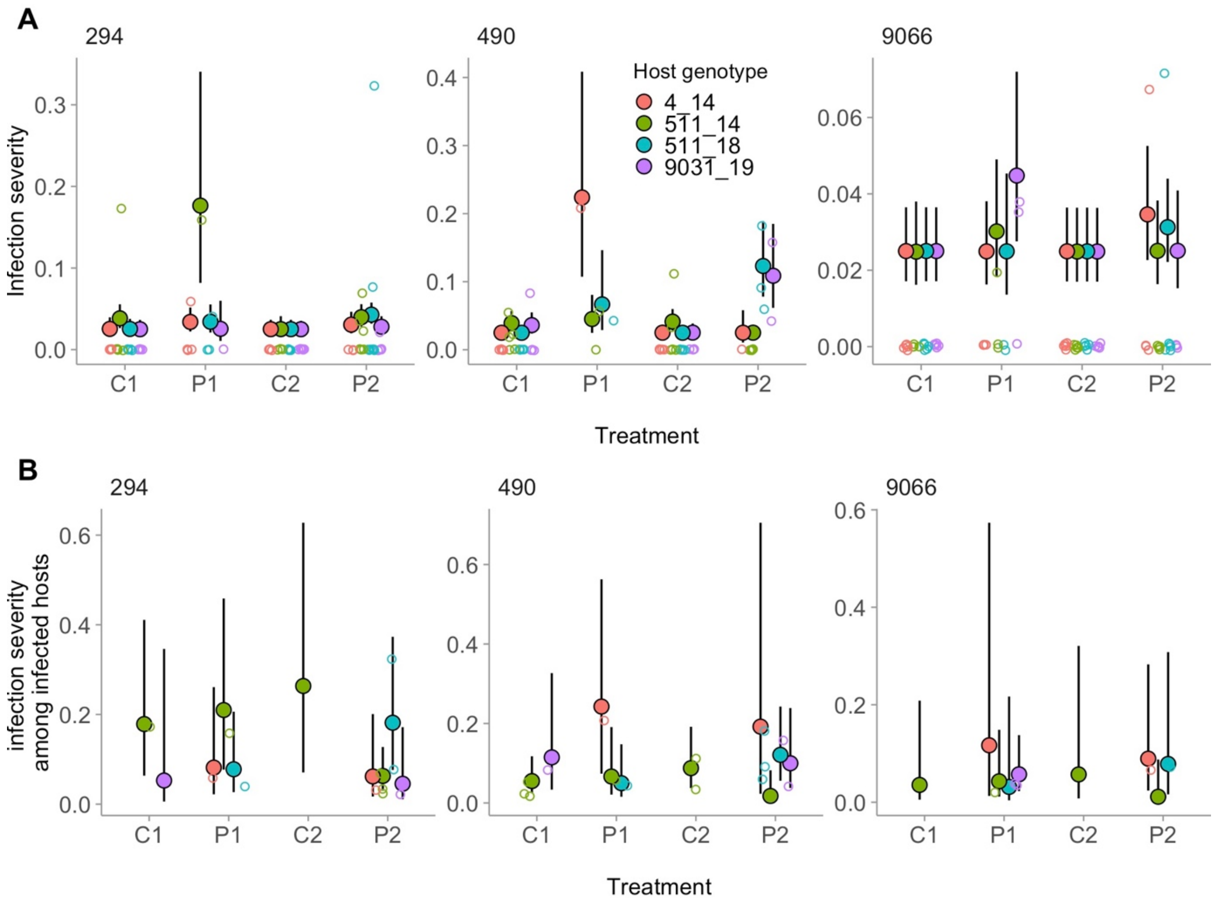


**Figure S1.** Results from the natural epidemic experiment. The effect of the priming treatment on a) infection severity; b) infection severity among infected hosts only. Plants were primed either 8 days (treatment P1) or 4 days (treatment P2) prior to being placed into the field. There were also C1 and C2 control plants set up at the same time (but mock inoculated). Filled points are model-estimated means, error bars are model-estimated 95% confidence intervals, and open points show the raw data. Panels are different host populations, and colors are different host genotypes. There was a significant three-way interaction in the model of infection severity. The reduced model of infection severity among infected hosts included significant two-way interactions between population and host genotype and between host genotype and experimental treatment.
